# Evaluation of a Dual Isolation Width Acquisition (DIWA) method for isobaric labelling ratio decompression

**DOI:** 10.1101/387878

**Authors:** Theodoros I. Roumeliotis, Hendrik Weisser, Jyoti S. Choudhary

## Abstract

Isobaric labelling is a highly precise approach for protein quantification. However, due to the isolation interference problem, isobaric tagging suffers from ratio underestimation at the MS2 level. The use of narrow isolation widths is a rational approach to alleviate the interference problem; however, this approach compromises proteome coverage. We reasoned that although a very narrow isolation window will result in loss of peptide fragment ions, the reporter ion signals will be retained for a significant portion of the spectra. Based on this assumption we have designed a Dual Isolation Width Acquisition (DIWA) method, in which each precursor is first fragmented with HCD using a standard isolation width for peptide identification and preliminary quantification, followed by a second MS2 HCD scan using a much narrower isolation width for the acquisition of quantitative spectra with reduced interference. We leverage the quantification obtained by the “narrow” scans to build linear regression models and apply these to decompress the fold-changes measured at the “standard” scans. We evaluate the DIWA approach using a nested two species/gene knockout TMT-6plex experimental design and discuss the perspectives of this approach.

## INTRODUCTION

Stable isotope labelling of peptides using isobaric reagents such as iTRAQ and TMT enables the multiplexed analysis of proteomes with deep quantitative coverage^1–2^. This barcoding strategy has provided comprehensive proteomic portraits of large collections of human cancer tissue samples^3–4^ and cell lines^5–6^, and has enabled the in-depth characterization of protein post-translational modifications in a quantitative fashion^7–10^. Isobaric tagging demonstrates high precision^11^ but imperfect accuracy due to ratio underestimation caused by co-fragmentation of ions with mass-to-charge ratios within the isolation window of the targeted precursors^12^. Although this problem rarely affects the direction of protein abundance change, many applications can significantly benefit from increased accuracy; examples include the determination of protein localization^13^, identification of specific protein-protein interactions in pull down assays^14–15^, and verification of protein deletions in gene knock-out or knock-down experiments or in samples with natural genomic variation^16^. Several groups have proposed solutions to alleviate the interference problem and thereby improve isobaric labelling quantification accuracy. Ow et al. demonstrated that the ratio compression can be decreased using high-resolution HILIC fractionation which achieves maximum orthogonality and reduces MS sampling complexity^17^. An alternative approach by Savitski et al. improves accuracy by fragmentation of the peptides close to their maximum chromatographic peak height^18^. Elimination of ratio distortion in a two-proteome model utilizing MS3 for further fragmentation of b- or y- ions specific to the targeted precursor was reported by Ting et al.^19^. McAlister et al. have further enhanced the sensitivity of the MS3 method using isolation waveforms with multiple frequency notches^20^. The latter is currently the method of choice for counteracting interference in isobaric labelling experiments using tribrid mass spectrometry. The QuantMode method developed by Wenger et al. is based on gas-phase purification, by manipulation of either mass or charge through expedient proton-transfer ion-ion reactions and has also been shown to improve quantitative accuracy^21^. Using data analysis methods, Wuhr et al. showed that precursor-specific quantitative information can be retrieved at the MS2 level from the complement reporter ion cluster^22^. Notably, Shliaha et al. evaluated the utility of ion mobility for additional precursor purification in data-dependent acquisition mode and presented evidence for improved accuracy, especially in combination with narrowed quadrupole isolation window^23^.

Prompted by empirical observations of isobaric-labelled peptide MS2 spectra, we argue that although a very narrow isolation window will result in severe loss of backbone fragment ions, rendering the spectra unsuitable for peptide identification, the reporter ion signals will remain intense enough to generate quantitative information for a significant portion of the spectra. Based on this assumption we have designed a Dual Isolation Width Acquisition (DIWA) method, in which each precursor is first fragmented with HCD using a standard isolation width for peptide identification and preliminary quantification, followed by a concomitant MS2 HCD fragmentation using a much narrower isolation width for the acquisition of quantification-only spectra with reduced interference. We leverage the quantitative values obtained by the “narrow” scans to build linear regression models and apply these to decompress the fold-changes measured at the “standard” scans. Here, we evaluate the DIWA method using a nested two species/gene knockout TMT-6plex model and discuss the potential of this approach.

## METHODS

### Experimental design overview

To alleviate the interference problem at the MS2 level we have designed an acquisition method on an LTQ-Orbitrap Velos in which each precursor is fragmented twice back-to-back with HCD using a different isolation width at each scan event (1^st^ MS2 scan 2.0 Th / 2^nd^ MS2 scan 0.1 or 0.2 or 0.3 Th) (Figure 1A). Slightly different collision energies are used in each of the two HCD spectra to mark their isolation width of origin. The aim of the method is to collect MS2 spectra devoid of interference in the isobaric tags which can be used to model the compression effect and to correct all quantitative values obtained at the standard isolation width. To evaluate this approach we have performed a TMT-6plex experiment using the mixed-proteome model^19^ in combination with the use of a CRISPR-cas9 gene knockout (KO). We analysed a mixture of Escherichia coli tryptic peptides at ratios 2:1, 4:1, 8:1 (×2 each) and tryptic peptides from three human cell lines (hiPSC ARID1A KO, hiPSC WT and CL-40) at ratios 1:1:1 (Figure 1B). Three TMT channels (129,130,131) were overlapping between E. coli and Human peptides whilst three channels (126,127,128) were used for interference-free E. coli peptides. To simulate lower-abundant changing proteins, we spiked the E. coli peptides at 2.5 to 20-fold lower total amount per channel compared to the human peptides. This represents a scenario of high interference; in a real experiment it is more likely that the differentially regulated proteins will cover the entire protein abundance dynamic range rather than the mid-to-low abundant portion only. The labelled peptides were fractionated with high-pH reversed-phase HPLC and the 25 fractions were analyzed on an LTQ Orbitrap Velos. The analysis of the fractions was performed three times at three different “narrow” isolation widths (0.1, 0.2 and 0.3 Th). Finally, linear regression models were generated by plotting the “standard” against the “narrow” isolation peptide logarithmic ratios for each sample comparison. All primary peptide log_2_ratios were calibrated through these models. We chose to model the compression effect by Deming regression, which is more appropriate when both the dependent and independent variables are measured with error, thus facilitating the comparison of two assays designed to measure the same analyte.

**Figure 1.**
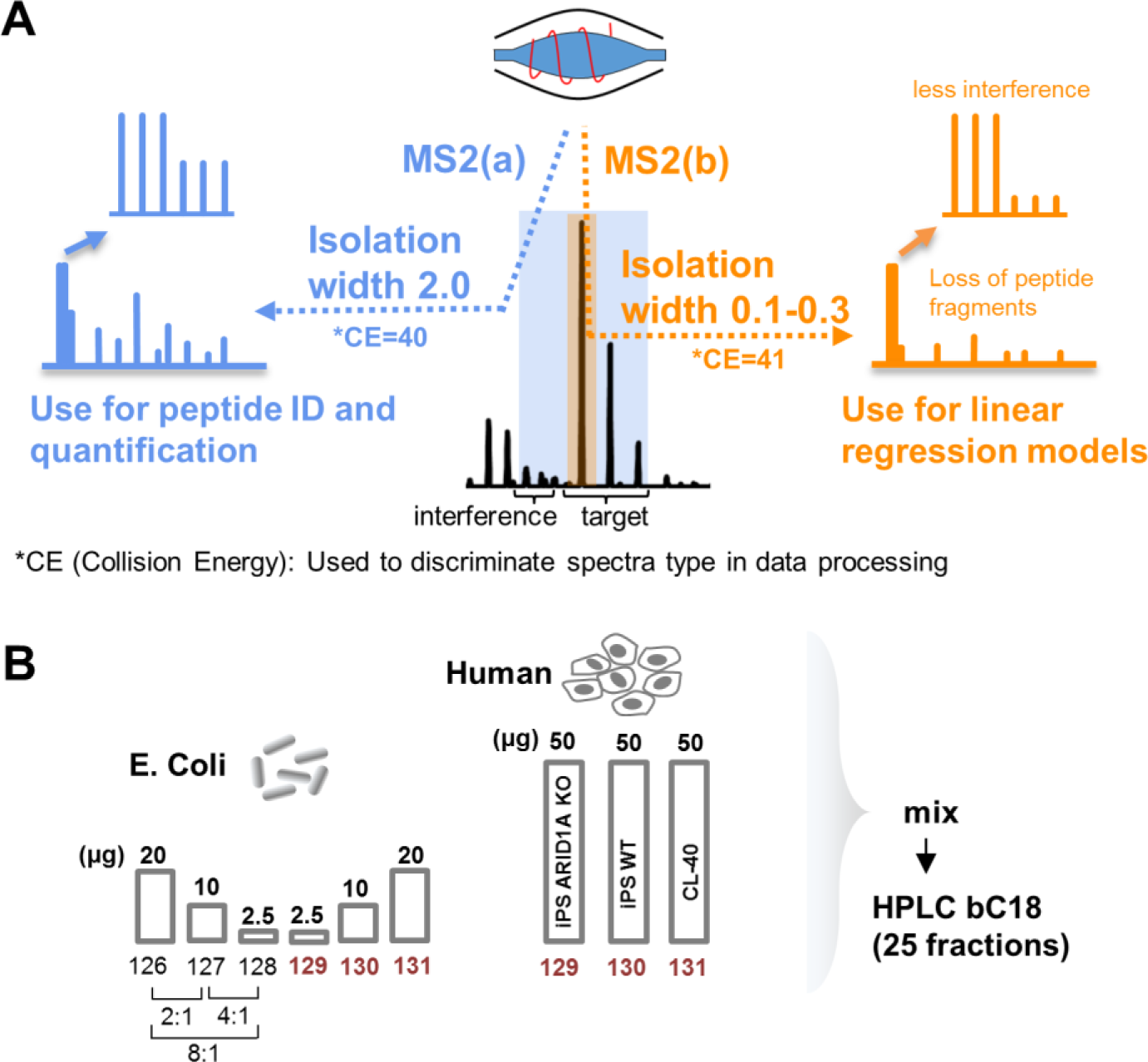
A) The DIWA method overview. B) Experimental design for the nested two species/gene knockout TMT-6plex model.

### Sample preparation and LC-MS analysis

The hiPSC CRISPR-cas9 and the CL-40 cell pellets were obtained as described previously^5^. The cell pellets were homogenized in 150 μL 0.1 M triethylammonium bicarbonate (TEAB), 1% sodium deoxycholate (SDC), 10% isopropanol with probe sonication for 3×5 sec with pulses of 1 sec at 40% amplitude (EpiShear) followed by boiling at 90 °C for 5 min. Samples were re-sonicated and centrifuged at 10,000 rpm for 10 min. Lyophilized Escherichia coli whole protein extract (Bio-Rad) was dissolved in 200 μL 0.1 M TEAB, 0.1% SDS divided into aliquots of 10 μL each and diluted up to 100 μL. The protein content of each aliquot was precipitated by the addition of 30 μL TCA 8 M for 30 min at 4 °C. The protein pellets were washed twice with ice-cold acetone and finally re-suspended in 40 μL 0.1 M TEAB, 0.1% SDS with probe sonication, before they were combined in a single E. coli pool. Protein concentration was measured with the Coomassie Plus Bradford Protein Assay (Pierce) according to manufacturer’s instructions. Duplicate aliquots of 20 μg (TMT: 126 and 131), 10 μg (TMT: 127 and 130), and 2.5 μg (TMT: 128 and 129) of E. coli protein and aliquots of 50 μg of hiPSC ARID1A KO (TMT: 129), hiPSC WT (TMT: 130) and CL-40 (TMT: 131) were prepared for trypsin digestion. Cysteines were reduced with 5 mM tris-2-carboxyethyl phosphine (TCEP) for 1 h at 60 °C and blocked by 10 mM iodoacetamide (IAA) for 30 min at room temperature in dark. Trypsin (Pierce, MS grade) solution was added at a final concentration of 70 ng/μL to each sample for overnight digestion. The peptide samples were finally labelled with the TMT-6plex reagents (Thermo Scientific) according to manufacturer’s instructions. The TMT peptide mixture was acidified with 1% formic acid and the precipitated SDC was removed by centrifugation. Offline peptide fractionation was based on high pH Reverse Phase (RP) chromatography using the Waters XBridge C18 column (2.1 x 150 mm, 3.5 μm, 120 Å) on a Dionex Ultimate 3000 HPLC system at a 0.85% gradient with flow rate 0.2 mL/min. Mobile phase A was 0.1% ammonium hydroxide and mobile phase B was 100% acetonitrile, 0.1% ammonium hydroxide.

LC-MS analysis was performed on the Dionex Ultimate 3000 UHPLC system coupled with the LTQ Orbitrap Velos mass spectrometer (Thermo Scientific). Samples were analysed with the Acclaim PepMap RSLC C18 capillary column (75 μm × 50 cm, 2 μm, 100 Å). Mobile phase A was 0.1% formic acid and mobile phase B was 80% acetonitrile, 0.1% formic acid. The gradient separation method was as follows: for 85 min gradient up to 38% B, for 10 min up to 95% B, for 10 min isocratic at 95% B, re-equilibration to 5% B in 5 min, for 10 min isocratic at 5% B. For the DIWA method, the five most abundant multiply charged precursors within 380-1500 *m/z* were selected with FT mass resolution of 30,000 and isolated for HCD fragmentation twice with isolation width 2.0 and 0.1 or 0.2 or 0.3 Th in an Nth order double play method (henceforth “standard” and “narrow” scans respectively). Normalized collision energy was set at 40 for the standard scans and at 41 for the narrow scans. Tandem mass spectra were acquired at 7,500 FT resolution with 40 seconds dynamic exclusion and 10 ppm mass tolerance. FT maximum ion time for full MS experiments was set at 200 ms and FT MSn maximum ion time was set at 50 ms. The AGC target vales were 3×10e6 for full FTMS and 5×10e5 for MSn FTMS.

### Data processing

The MS2 spectra collected with collision energy 40 (“standard” scans) were searched against Uniprot human (reviewed only) and Escherichia coli entries using SequestHT in Proteome Discoverer 2.2. The precursor mass tolerance was set at 30 ppm and the fragment ion mass tolerance was set at 0.02 Da. Static modifications were: TMT6plex at N-termini/K, and carbamidomethyl at C. Dynamic modifications included oxidation of M and deamidation of N/Q. The data were processed twice, first with maximum collision energy 40 for quantification using the “standard” scans and second with minimum collision energy 41 for quantification using the “narrow” scans (Reporter Ions Quantifier node). Quantification was based on un-normalized signal-to-noise (S/N) values. Peptide confidence was estimated with the Percolator node. Peptide FDR was set at 0.01 and validation was based on q-value and decoy database search. Deming regression was performed in RStudio with the “deming” package. A matrix containing the log_2_ratios: 131/129, 131/130 and 130/129 for the standard (IW2) and narrow (IW01) scans was used as input. The R code and the regression models are provided in Supporting information. The mass spectrometry proteomics data have been deposited to the ProteomeXchange Consortium via the PRIDE^24^ partner repository with the dataset identifier PXD010571.

## RESULTS

To evaluate the depth of proteome coverage and the accuracy level that can be achieved by the dual isolation width acquisition method, we performed a proof-of-concept TMT6plex-based analysis of varying amounts of spike-in E. coli protein extract into human protein lysates of equal total amounts representing different cell types. For straightforward implementation of the method, we used an LTQ-Orbitrap Velos platform in which the method editor already allows the setup of dual acquisitions (e.g. CID-HCD). The application of the DIWA method in the two-proteome model resulted in the quantification of 6,724 total human and E. coli unique protein groups at isolation width (IW) 2.0. This level of proteome coverage is in line with previous studies using similar instrumentation^9^. A subset of 6,132 (91%) proteins were also fully quantified at the narrowest isolation width 0.1, confirming that the isobaric tags remain at detectable levels for the majority of the peptides despite the significant loss of the peptide fragment ion signals (Supplemental Table 1). As expected, the percentages of quantified PSMs, peptides and protein groups increased as the narrow scan windows were opened to 0.2 and 0.3 (Table 1, Supplemental Tables 2 and 3). The overall lower quantification coverage of E. coli peptides and proteins is due to their lower abundances compared to human proteins and due to the 8-fold lower sample load in two of the TMT channels. These low-intensity TMT channels were more frequently below the quantification limit compared to the human samples, resulting in smaller number of fully quantified E. coli PSMs. Notably, quantification at narrow IW was more efficient for doubly charged peptides (Supplemental Figure 1A) with medium to high precursor intensity (Supplemental Figure 1B). The charge state dependency is possibly due to the fact that the monoisotopic peak of doubly charged peptides is the most intense within their isotopic cluster yielding more efficient isolation. However, multiply charged peptides appeared to have overall lower precursor intensities (Supplemental Figure 1C) suggesting that isolation efficiency depends on both the isotopic cluster pattern and precursor intensity.

**Table 1.**
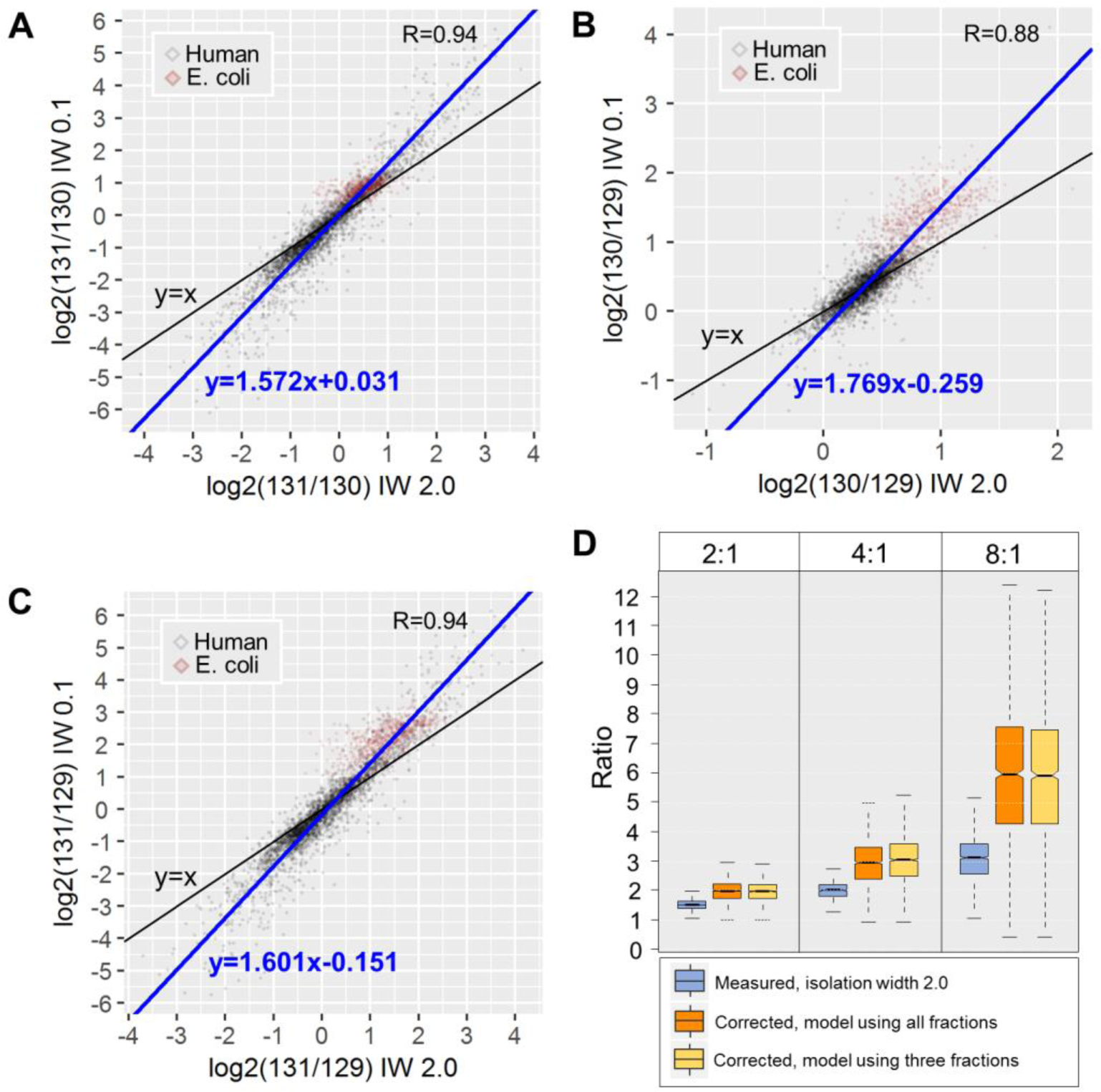
Number and comparative percentages of PSMs, unique peptides and protein groups quantified at each isolation width for human and E.coli samples.

Next, we evaluated the correction efficiency of the DIWA method using the E. coli PSMs fully quantified in the different isolation widths. Significantly less TMT signal distortion was found using the narrow isolation widths as shown by the difference between the medians of the expected (without interference) and measured (with interference) scaled quantitative values between the replicate channels (Figure 2A). Although the compression effect was not eliminated, the narrow isolations yielded significantly improved ratios (Figure 2B). Specifically, in isolation width 0.1, the percent error was decreased from 59% to 35% for the higher ratio 8:1 and from 21% to 11% for the lower ratio 2:1. For example, while 67% of the ratios measured at isolation width 2.0 were below 4 for the expected ratio 8:1, 73% of these were above the 4-fold threshold using the narrow acquisition. Additionally, the ratio compression effect showed a linear correlation (R^2^=0.99) with the isolation width; as found by the median ratios at each one of the narrow acquisitions (Figure 2C). This suggests that the compression effect could be modelled by the isolation width gradient to predict the ratios at isolation widths close to zero. Because the use of the narrowest isolation width yielded the smallest interference, for all downstream analysis we use only the data obtained at isolation width 0.1. Two example identification and quantification spectra matched to ghrB (E. coli) and ARID1A (human) peptides are shown in Figure 3. Both peptides suffered significant ratio compression at the “standard” isolation width due to high precursor interference, however the second HCD scan at isolation width 0.1 provided more accurate quantification (Figure 3, right panels). Specifically, at the “standard” scan, the ARID1A peptide displayed a 3.9-fold down-regulation in the ARID1A knock-out cells suggesting an in-complete silencing of the gene. However, the narrow MS2 scan of the same precursor ion revealed a 19-fold reduction of the protein product. The latter is more likely to reflect a complete knock-out, particularly when the previously described spatial constraints on protein quantification by TMT^25^, that do not permit proper discrimination of truly missing proteins are taken into consideration.

**Figure 2.**
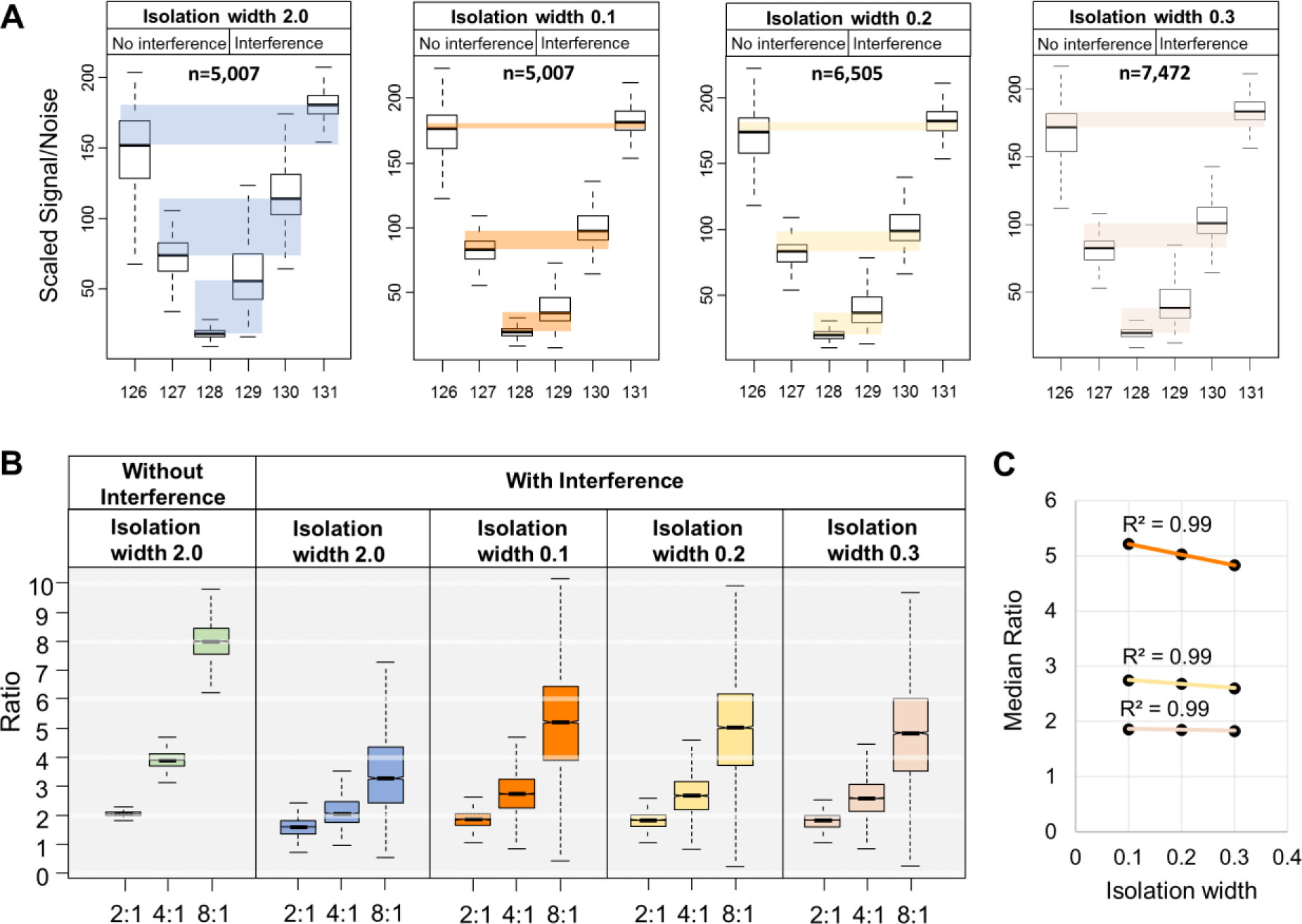
Reduction of interference by the DIWA method. A) Box plots of the scaled TMT signal-to-noise values of E. coli PSMs across the six TMT channels at the different isolation widths. The embedded colored boxes highlight the difference of the medians between the expected (No interference) and measured (Interference) values for each replicate pair. B) Box plots of the E. coli PSM ratios for each theoretical ratio at the different isolation widths without and with interference. C) Scatter plot of the isolation width setting (x-axis) against the median ratio (y-axis) for each theoretical ratio.

**Figure 3.**
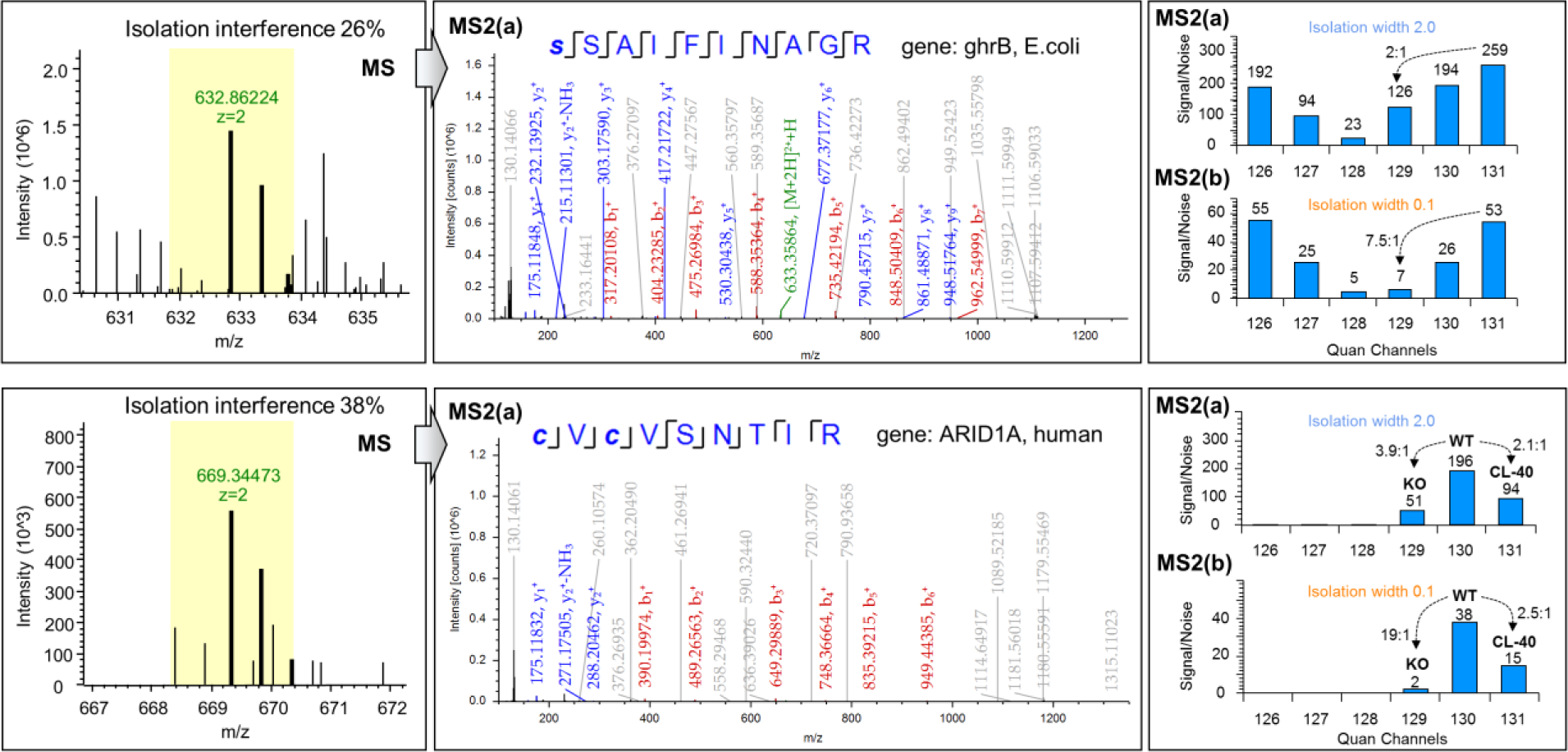
Example spectra for E. coli and human peptides. Precursor ion cluster, annotated HCD MS2 fragment ion spectrum and TMT signal-to-noise values at “standard” and “narrow” isolation widths for a peptide matched to E. coli ghrB gene (top panel) and for a peptide matched to human ARID1A (bottom panel). Unassigned peaks are shown in grey font on the MS2 spectra.

As the narrow isolation widths did not provide quantification for about 25-35% of the respective “standard” scans, we next aimed to model the compression effect using linear regression and to calibrate all primary quantifications obtained at the “standard” scans. Upon manual examination of the quantification spectra, we found that the narrow isolation widths were not always effective in reducing isolation interference. To identify the E. coli spectra with significant reduction of the interference upon the application of narrower precursor selection and therefore to model the compression effect more accurately, we computed the ratio 129(IW 0.1)/129(IW 2.0) of the scaled abundances as a metric for the magnitude of correction. In this instance, a low ratio (large difference in signal intensity) would suggest effective correction by the narrow scan whereas a high ratio would suggest insufficient correction or spectra with originally low interference. Examples of quantification spectra with low and high 129(IW 0.1)/129(IW 2.0) ratio are shown in Supplemental Figure 2A. To identify which peptide features are associated with effective correction by the narrow scans, we next correlated this ratio metric with peptide m/z, charge, precursor intensity and isolation interference (the percentage of ion signal not attributed to the targeted precursor within a specified isolation window as reported by ProteomeDiscoverer software) (Supplemental Figure 2B). We found that peptides with lower m/z, charge and precursor intensity (positively correlated) and high isolation interference (negatively correlated) are more effectively corrected by the narrow scans. Therefore, we can enrich for peptides that are effectively corrected by the narrow scans in samples with unknown protein abundances, by applying cutoffs to these features. Consequently, we selected doubly charged PSMs, with m/z and precursor intensity smaller than the median of all PSMs (<698.4 and <2.4E+6 respectively) as well as isolation interference greater than the median of all PSMs (>18.2%) as input for the Deming regression analysis (n=3,616). To model the compression effects we generated the scatterplots of the selected PSMs using the logarithmic ratios from IW 2.0 scans against their counterparts at IW 0.1. We observed a linear response with Pearson’s R>0.88 and Deming regression slope>1.57 for three comparisons (131/130, 130/129 and 131/129) indicative of the compression effect (Figure 4A, 4B and 4C). Using the slope and the intercept of these linear models we calibrated the log_2_ratios acquired at IW 2.0 for all human and E. coli PSMs. To evaluate the decompression efficiency, we retrieved the calibrated E. coli PSM log_2_ratios, converted to 2^log_2_ratio^ and computed the mean ratio per protein. This analysis showed that the regression-based calibration could decompress the original ratios up to 1.9-fold on average (Figure 4D). For example, in the expected ratio 8:1 only 28 proteins (2%) had a ratio greater than 5 at the standard isolation width however in the calibrated values, 820 proteins (66%) were above this threshold. This improvement can have important implications in statistical analysis and identification of differentially expressed proteins when specific cut-offs are applied. As the dual acquisition method is associated with longer cycle times, we tested whether efficient predictive models could be built using a smaller subset of PSMs from only three randomly selected fractions. Indeed, a very similar degree of decompression could be achieved from only a subset of the fractions (Figure 4D, Supporting information). This suggests that all peptide fractions could be analysed with a usual TMT method for maximum proteome coverage, followed by the DIWA re-run of only a few fractions for regression analysis and retrospective decompression of all the original ratios. Additionally, correlation analysis showed that the percent error at IW 2.0 is positively correlated with charge state and isolation interference and negatively correlated with precursor intensity, m/z and Sequest cross-correlation score (Xcorr) (Supplemental Figure 2C). These characteristic features could be utilized to build more accurate predictive models using machine learning approaches (e.g. Support Vector Regression). Overall, our feasibility experiment shows that the acquisition of an additional HCD scan event at a narrow isolation width immediately after the acquisition of a standard MS2 scan for the same precursor, can be used to enhance accuracy of quantification for the majority of the identified peptides. Moreover, the dual isolation data can be used to model the compression effect by linear regression extending the coverage of the ratio decompression.

**Figure 4.**
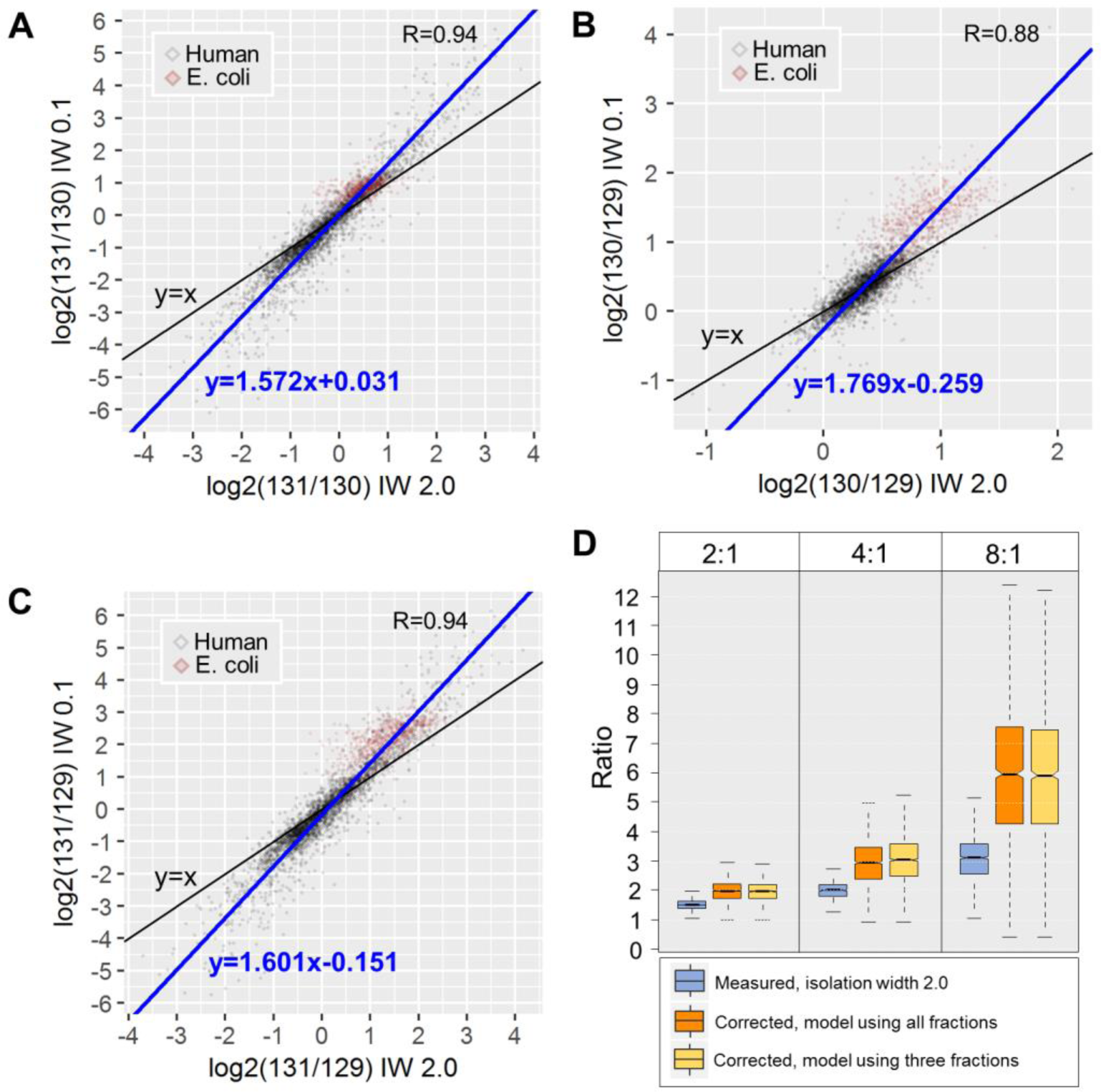
Deming regression models and ratio decompression. Scatter plots of the (A) 131/130, (B) 130/129 and (C) 131/129 logarithmic ratios at isolation width 2.0 (x-axis) versus isolation width 0.1 (y-axis). D) Box plots of the E. coli proteins ratios (average of PSM ratios) before and after decompression at the different theoretical ratios.

## CONCLUSIONS

The selection of precursor ions using narrow isolation windows is a rational approach to reduce peptide co-fragmentation and therefore to improve isobaric labelling-based quantification at the MS2 level. However, this approach yields very low proteome coverage as the narrow precursor selection results in poor peptide fragment ion spectra. To address this problem, we have designed a novel method based on sequential HCD-HCD activation in a dual isolation width mode followed by modelling of ratio compression and correction. We have tested the method on an LTQ Orbitrap Velos system with a common Nth-order double play method. Importantly, apart from some additional data analysis steps, the DIWA approach does not require major changes in sample preparation protocols or specialized instrument configuration adjustments of the LTQ-Orbitrap systems (Velos and Elite). Using a two-proteome model and a CRISPR-cas9 gene knockout, we show that the method achieves comprehensive proteome coverage and preliminary quantification of all peptides while the additional narrow isolation width can improve the quantitative accuracy for a significant portion of these. Furthermore, the low-interference spectra can be used as “pseudo-internal standards” to model the compression effect by linear regression in a sample-specific manner. The enhancement offered by the DIWA method is anticipated to be universal to other platforms, such as benchtop Q-Exactives or Q-TOFs, that can only perform isobaric labelling quantification at the MS2 level. Moreover, the combination of DIWA with previously described approaches such as gas-phase purification and ion mobility could further improve the accuracy of isobaric labelling approach at the MS2 level. Future developments in mass spectrometry technology, which improve isolation efficiency and analytical speed in combination with intelligent precursor selection decision trees could further boost the sensitivity and accuracy of the method. We conclude that the DIWA approach can provide significant ratio decompression in isobaric labelling at the MS2 level, and that the current implementation offers the foundations for further developments and offers universal applicability.

## AUTHOR CONTRIBUTIONS

Conceptualization: T.I.R., J.S.C.; Experiments: T.I.R.; Data analysis: T.I.R., H.W.; Writing original draft: T.I.R., J.S.C.; Final draft/editing: all

## NOTES

The authors declare no competing financial interest.

## ACKNOWLEDGMENTS

We thank Ultan McDermott and Stacey Price for donating the CL-40 cell pellet, David J. Adams for donating the human iPS WT and ARID1A KO cell pellets and Daniel Bode for discussions about data analysis. This work was funded by a core grant from the Wellcome Trust (098051).

## Abbreviations

DIWA: Dual Isolation Width Acquisition
HCD: Higher-energy Collisional Dissociation
HILIC: Hydrophilic Interaction Chromatography
iTRAQ: Isobaric Tags for Relative and Absolute Quantification
IW: Isolation Width
KO: Knockout
PSM: Peptide Spectrum Matches
RP: Reversed Phase
TMT: Tandem Mass Tags

